# Identification of the EH CRISPR-Cas9 system on a metagenome and its application to genome engineering

**DOI:** 10.1101/2022.10.31.514646

**Authors:** Belen Esquerra-Ruvira, Ignacio Baquedano, Raul Ruiz, Almudena Fernandez, Lluis Montoliu, Francisco JM Mojica

## Abstract

Non-coding RNAs (crRNAs) produced from clustered regularly interspaced short palindromic repeats (CRISPR) loci, and CRISPR associated (Cas) proteins of the prokaryotic CRISPR-Cas systems, form complexes that interfere with the spread of transmissible genetic elements through Cas-catalysed cleavage of foreign genetic material matching the guide crRNA sequences. The easily programmable targeting of nucleic acids enabled by these ribonucleoproteins has facilitated the implementation of CRISPR-based molecular biology tools for *in vivo* and *in vitro* modification of DNA and RNA targets. Despite the diversity of DNA-targeting Cas nucleases so far identified, native and engineered derivatives of the *Streptococcus pyogenes* SpCas9 are the most used for genome engineering, at least in part due to its catalytic robustness and the requirement of an exceptionally short motif (5’-NGG-3’ PAM) flanking the target sequence. However, the large size of the SpCas9 variants impairs the delivery of the tool to eukaryotic cells and smaller alternatives are demanded. Here we identify in a metagenome a new CRISPR-Cas9 system associated with a smaller Cas9 protein (EHCas9) that targets DNA sequences flanked by 5’-NGG-3’ PAMs. We develop a simplified EHCas9 tool that specifically cleaves DNA targets and is functional for genome editing applications in prokaryotes and eukaryotic cells.

## Introduction

CRISPR-Cas components have been identified in genetic elements of many bacteria and most archaea (https://crispr.i2bc.paris-saclay.fr/crispr/). They play diverse roles based on CRISPR RNA-guided nucleic acid-targeting (either DNA or RNA targets, depending on the system) by surveillance nucleoprotein complexes that involve effector Cas proteins bound to the guide [1,2]. Typically, recognition by the surveillance complex of a guide-complementary sequence [3] in infectious genetic elements such as viruses [4] or plasmids [5] results in target cleavage [6] catalysed by a Cas nuclease, CRISPR-Cas functioning in this case as a defence system [7]. The current classification of these systems distinguishes two classes, six types and several subtypes [8–10]. Class 2 systems (including types II, V and VI) mainly differ from class 1 in the composition of the surveillance complex which contains a single effector Cas protein as opposed to the multiprotein complex characteristic of class 1. Type II systems are categorized into three subtypes (II-A, II-B, and II-C), sharing the signature effector protein Cas9 [8] that has two DNA nuclease domains, RuvC and HNH [11]. Also, in contrast with class 1 systems that are guided by just a noncoding CRISPR RNA (crRNA) [3], Cas9 is guided by a hybrid RNA (dual gRNA) composed of crRNA and a transactivating crRNA (tracrRNA) [12,13]. Each mature crRNA comprises partial sequences of one repeat and the adjacent repeat-intervening region (named spacer) resulting from cleavage at the repeats in CRISPR RNA transcripts (pre-crRNA) typically transcribed from a promoter in the so-called leader sequence flanking the CRISPR array, and further 5’-trimming. In addition to Cas9 and tracrRNA, non-Cas RNases (*i.e*., RNase III in some subtypes) may be required or not (depending on the system variant) for crRNA generation [12–15]. The tracrRNAs are usually encoded in the vicinity of the CRISPR-*cas* loci and contain an anti-repeat region (the anti-repeat base pairs with the repeat in CRISPR RNA molecules) and partially palindromic sequences that adopt stable stem-loop structures to bind Cas9 [12,16]. The Cas9 in the surveillance complex scans dsDNA for short sequences called protospacer adjacent motifs (PAMs) [17–19]. When a compatible PAM is bound by the PAM-Interacting (PI) domain of Cas9, the DNA is locally destabilized facilitating interrogation of the upstream sequence for complementarity with the spacer region of the crRNA [20]. Base pairing between the gRNA and the spacer-complementary sequence in the target will form an R-loop, relocating the PAM-proximal region of the two DNA strands at the RuvC and HNH catalytic sites to produce a double-strand break at a fixed distance from the PAM [6,13,20–22]. The easy programming of effector Cas proteins for binding to nucleic acids has led to the implementation of diverse CRISPR tools for disease diagnosis [23,24] and genome modification in heterologous hosts. Such applications include genome editing in a wide range of organisms, from bacteria [25,26] to human eukaryotic cells [27–29], and the production of sequence-specific antimicrobials [30]. Primarily, the cleavage features (*i.e*., specific single double-strand break) and the simplicity of class 2 surveillance complexes in terms of the number of proteins required, have made DNA-targeting class 2 systems (i.e., type V and type II) the gold standard of CRISPR technology [31]. However, the native type II systems have constraints in this regard related to cell delivery of the Cas9 tool due to the large size of the nucleoprotein complex and the requirement of specific PAM sequences for efficient target recognition [32]. Cas9:gRNA tools have been simplified by linking shortened tracrRNA and crRNA molecules in a single-guide RNA (sgRNA) [13]. Among the Cas9 orthologs so far identified and biochemically characterized [33–36] the subtype II-A Cas9 from *Streptococcus pyogenes* (SpCas9) is the most widely used constituent of type II CRISPR-based tools [37]. Moreover, reconstructed Cas9 ancestors [38] and laboratory evolved [39,40] or engineered [33–36,41–43] Cas9 variants with smaller sizes and different PAM requirements have been harnessed to the CRISPR technology. Yet, the search for Cas9 orthologs in prokaryotic genomes [44–46] and metagenomes [47,48] is expanding the catalogue of CRISPR tools and, allegedly, will continue to provide new alternatives to circumvent their limitations.

Here we characterized a new type II-C system, referred to as the EH CRISPR-Cas system, identified in a metagenome dataset from a water sample of a natural environment in Spain. Compared to previously reported type II nucleases, the associated Cas9 (EHCas9) features size in the range of the smallest orthologues and have distinct PAM preferences. Based on this system, we have developed an EHCas9 tool that functions both in eukaryotic and prokaryotic cells and can be used for genomic engineering.

## Results

### Detection of type II CRISPR-Cas systems in El Hondo Natural Park

To discover new type II CRISPR-Cas systems we explored microbial aquatic communities from El Hondo Natural Park, in Spain. A water sample was sequentially filtered to obtain a subcellular fraction, hopefully containing streamlined *cas9* genes. A megabase-scale metagenomic dataset generated from this sample was explored to detect Cas9-encoding CRISPR-Cas systems. We selected for further analysis a CRISPR-Cas II system (here named EH CRISPR-Cas) that was tentatively assigned to the II-C subtype considering the composition of the *cas* locus, made of the adaptation genes *cas1* and *cas2* preceded by the effector *cas9* (*ehcas9*) gene, and a CRISPR array consisting of two 36-bp repeats separated by a 29-bp spacer (Fig. 1). This might be a truncated array as it is located at the end of the contig and assemblers often collapse at repetitive regions [49,50]. A putative tracrRNA-encoding gene was identified upstream of *ehcas9* as an ~100-bp region, flanked by a predicted promoter and Rho-independent terminator motifs, containing an anti-repeat sequence partially complementary to the associated CRISPR units. So that the tracrRNA could base pair with the cognate crRNA, transcription of the tracrRNA gene from its anticipated promoter implies that the associated CRISPR array should be transcribed in the same direction, either from a leader located at the array flank missed in the contig or, as documented for other type II-C systems [14,51] from promoters in the repeats.

**Fig. 1.**
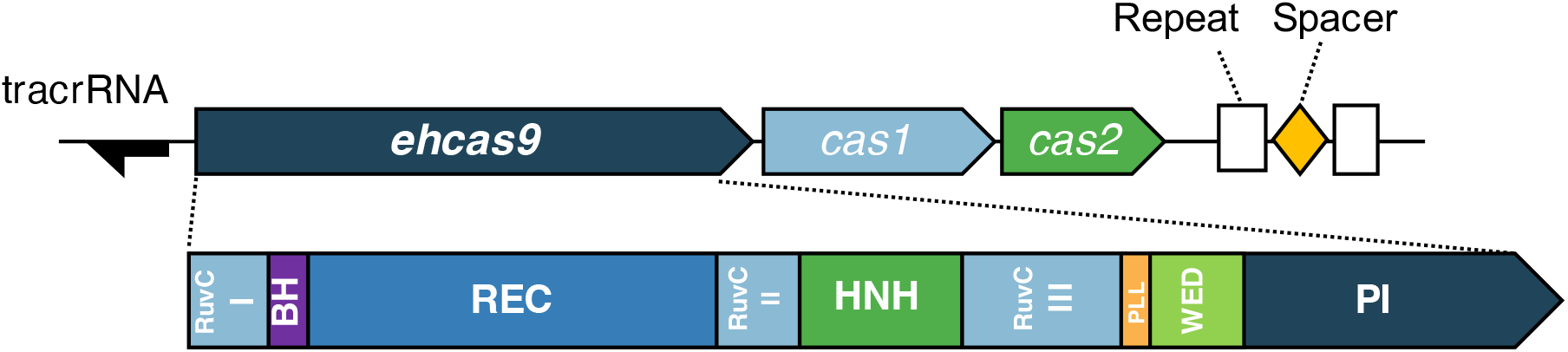
Schematic representation of the EH CRISPR-Cas locus and domains of the associated Cas9 protein. The EH CRISPR-Cas locus comprises three *cas* genes in the order of *cas9* (denoted *ehcas9*) - *cas1* - *cas2* (boxes pointing toward the direction of transcription), and two 36-bp CRISPR units (Repeat, white rectangles) separated by a 29-bp spacer (diamond). The location of a putative tracrRNA gene is represented as a black arrow pointing towards the predicted direction of transcription. The EHCas9 protein domains (depicted below the locus representation) inferred from the alignment with orthologues (Supplementary Figure S2), include the RuvC (I, II and III motifs), Bridge Helix (BH), recognition (REC), HNH, Phosphate Lock Loop (PLL), WED and PAM-Interacting (PI) domains.

The *ehcas9* ORF would produce a 1070 aa protein (termed EHCas9) of ca.120 kDa. Comparison of the EH protein sequence using the BLASTp tool, disclosed amino acid identity < 68% with Cas9 proteins in the NCBI non-redundant protein sequences database. Multiple sequence alignments with structurally characterized Cas9 orthologs (Fig. S2) revealed the typical domain organization of the protein family (Fig. 1) [21,35,36,52–54] and 3D structure prediction (Figures S3A and S3B) confirmed that EH protein adopts the family characteristic bi-lobed architecture, formed by REC and NUC lobes. Moreover, EHCas9 features conserved catalytic residues in both the RuvC (D11, E521, H747 and D750) and HNH (D605, H606 and N629) nuclease domains (Fig. S2). In contrast, as previously observed for most Cas9 proteins [36], the sequence of the PI domain diverges considerably from that of other orthologues. This distinctiveness was corroborated by comparison between the predicted 3D structure of EHCas9 and solved structures of Cas9 proteins. Figures S3C and S3D illustrate the differences between the PI domains of CdCas9 and EHCas9 while maintaining the conserved core fold of anti-parallel β-sheets. Collectively, these observations suggested that EHCas9 might act as a crRNA:tracrRNA-guided nuclease similarly to biochemically characterized orthologues but recognizing distinct PAMs.

The evolutionary relationship of EHCas9 was analysed by reconstructing a phylogenetic tree including 798 Cas9 amino acidic sequences (Fig. 2). The topology of the tree was consistent with the ten Cas9 clades reported by Gasiunas and collaborators [46]. EHCas9 falls within clade IX of the II-C subtype and is distantly related to Cas9 proteins commonly used in genome editing, being Cas9 from *Sulfitobacter donghicola* the most closely related among biochemically characterized orthologues.

**Fig. 2.**
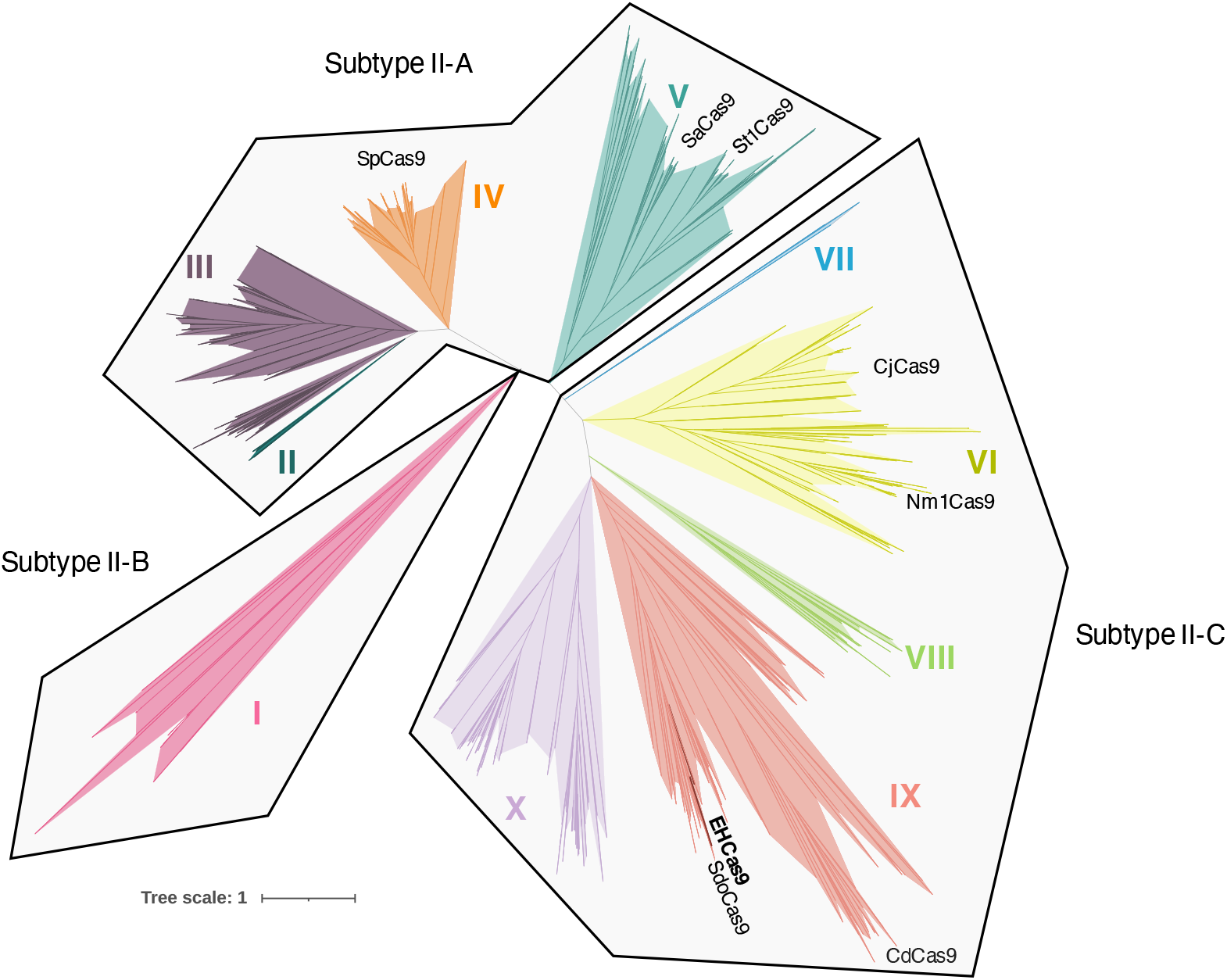
Evolutionary relationship of EHCas9. Phylogenetic tree of EHCas9 and 798 orthologues. Clades II (dark green), III (purple), IV (orange) and V (dark blue) belong to subtype II-A, clade I (pink) to subtype II-B, and clades VI (yellow), VII (blue), VIII (green), IX (red) and X (lilac) to subtype II-C. Cas9 of *Sulfitobacter donghicola* (SdoCas9) and orthologues commonly used for genome editing (SaCas9: *Staphylococcus aureus* Cas9; SpCas9: *Streptococcus pyogenes* Cas9; NmCas9: *Neisseria meningitidis* Cas9; CjCas9: *Campylobacter jejuni* Cas9; CdCas9: *Corynebacterium diphteriae* Cas9; StCas9: *Streptococcus thermophilus* Cas9) are labelled at their approximate position in the tree.

### Guide and PAM requirements for EHCas9-mediated DNA cleavage

To check *in vivo* DNA cleavage activity of EHCas9 guided by the predicted tracrRNA and a cognate crRNA, we inserted into *E. coli* plasmids the putative EH tracrRNA-encoding sequence and the *ehcas9* gene under inducible promoters, along with a minimal CRISPR array, composed of one spacer flanked by two EH repeats, transcribed from a constitutive promoter. We expected that, as shown for other type II systems [12,51,55,56], the pre-crRNA would be properly processed in the heterologous *E. coli* host to generate mature, guiding-proficient crRNAs. Then, we performed transformation assays of cells carrying the three EH CRISPR-Cas components, with a set of pSEVA431-derivative (spectinomycin resistance) plasmids containing a target sequence flanked from the 3’-end of the spacer-matching strand (i.e., the PAM region) by different sequences. Given the tolerance for any nucleotide in the first position of the PAM region exhibited by most Cas9 proteins [46], thymine was kept invariable in this location and random nucleotides at the 2nd, the 3rd, and the 4th positions (consensus 5’-TNNN-3’) were tested (Table S5). Analogous experiments were performed without the *ehcas9* gene. Spectinomycin resistant colonies were collected from each transformation experiment and PCR amplified DNA fragments including the PAM region were HTS sequenced. The comparison of the incidence of each sequence at the PAM region in the presence or absence of EHCas9 revealed that guanine was underrepresented in the 2nd and the 3rd positions when the protein was produced (Fig. 3A), but no difference in the frequency of any specific nucleotide was observed in the 4th position. These results demonstrate that EHCas9 can elicit specific interference against target plasmids if there is a guanine in the 2nd and the 3rd positions of the PAM. They also support EH tracrRNA identity alongside the inferred direction of transcription of the associated CRISPR array. Furthermore, they prove that a guiding-proficient crRNA is generated in *E. coli* from the engineered EH pre-crRNA under the conditions assayed.

**Fig. 3.**
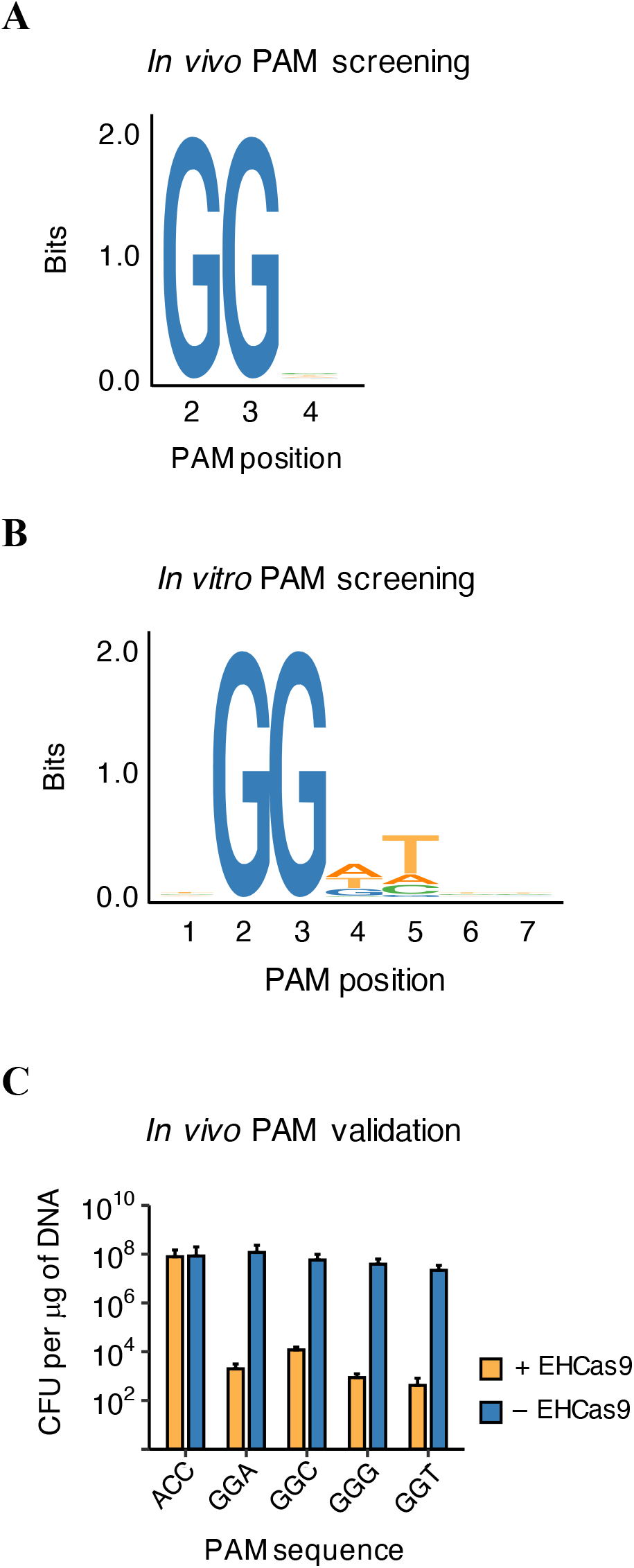
EHCas9 PAM screening and validation. (**A**) Sequence logo of the PAM region preferred by EHCas9 for target cleavage, as determined by *in vivo* screening of a PAM library. Nucleotide positions 3’ from the end of the spacer-matching strand of the target are indicated. Nucleotides from the 2nd to the 4th position were tested (the first position was kept invariable, corresponding to thymine). (**B**) Sequence logo of the consensus PAM preferred by EHCas9 for target cleavage as determined through *in vitro* screening. Nucleotide positions 3’ from the end of the spacer-matching strand of the target are indicated. Nucleotides from the 1st to the 7th position were tested. (**C**) *In vivo* PAM validation. The efficiency of transformation (number of colony-forming units – CFU - per mg of plasmid DNA) of *E. coli* cells expressing (+ EHCas9) or not (-EHCas9) EHCas9 in addition to a guide EH crRNA and the predicted EH tracrRNA, with plasmids carrying a target adjacent to sequences varying in the 2^nd^, the 3^rd^ and the 4^th^ positions (ACC, GGA, GGC, GGG, GGT) of the PAM region, is represented. Data are the mean of three replicates (error bars correspond to the standard deviation).

For the implementation of a simplified EHCas9 RNA-programmable DNA digestion tool, a single guide RNA (EH sgRNA) was inferred from the biochemically validated sequence of the type II-C system in *S. donghicola* (Fig. 2). After comparing the crRNA and tracrRNA of the two systems, we conceived a 118-nt long EH sgRNA, composed of a 23-nt customizable spacer region and a 95-nt long constant sequence (sgRNA backbone) consisting of an 18-nt truncated repeat, a 4-nt linker (tetraloop 5’-GAAA-3’) and a 73-nt EH tracrRNA fragment containing the anti-repeat followed by a sequence presumably adopting two stem-loop structures (Fig. 4).

**Fig. 4.**
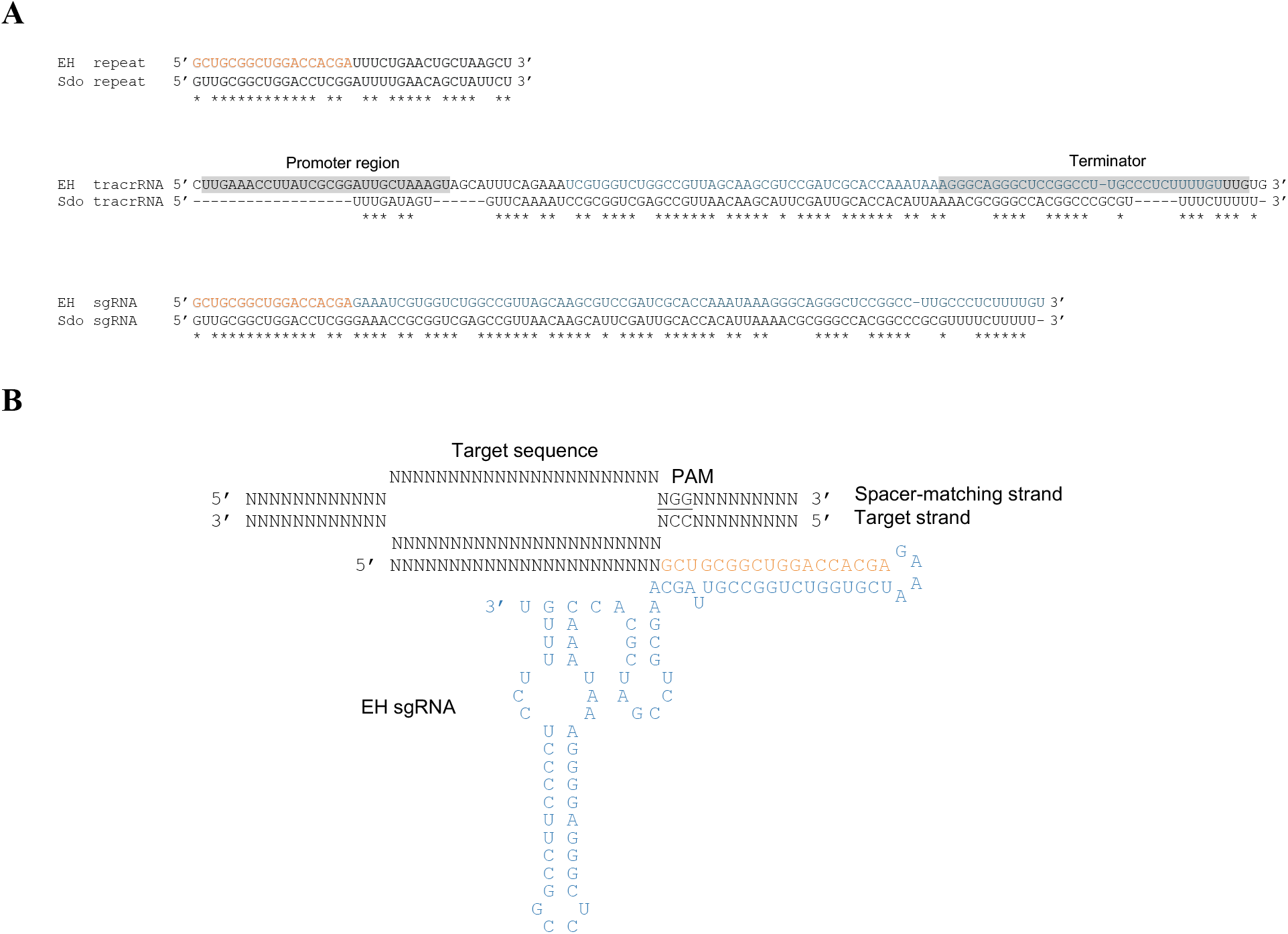
EH sgRNA design. (**A**) RNA sequence alignments of the EH repeat, the predicted EH tracrRNA gene region and the designed EH sgRNA with the *S. donghicola* (Sdo) repeat, tracrRNA coding sequence and sgRNA, correspondingly. Matching positions are marked with asterisks. Promoter and terminator regions predicted for the EH tracrRNA gene are grey shaded. The constant region of the EH sgRNA sequence was conceived by joining the EH repeat sequence in orange and the EH tracrRNA sequence in blue. (**B**) Scheme of the EH sgRNA including a generic 23-nt spacer base-paired to the target strand in a DNA substrate containing a spacer-matching sequence and a compatible PAM (underlined). The EH sgRNA anti-repeat and tracrRNA sequences comprising the linker (tetraloop 5’-GAAA-3’), the anti-repeat and the two stem-loop forming segments, are coloured as in panel A.

To test the functionality of the EH sgRNA and expand the PAM inferred from the *in vivo* screening, the first seven positions of the PAM region were tested using the cell-free *in vitro* translation (IVT) procedure as previously described [46]. This PAM screening was carried out in collaboration with CasZyme company, using EHCas9 and an *in vitro* transcribed EH sgRNA targeting a 7-nt PAM randomized plasmid library (Table S5). MgCl_2_ was included in the reaction as it has been shown that divalent cations are required by Cas9 proteins to adopt the cleavage-competent state [13,57–59]. Sequence analysis revealed target cleavage, corroborating the functionality of the engineered EH sgRNA. In line with most previously characterized Cas9 nucleases [13,46], cleavage was preferentially observed between nucleotides at the 3rd and 4th positions from the PAM on both strands of the target, suggesting blunt-end formation. The analysis of the PAM region (Fig. 3B) confirmed that, in agreement with the results of the *in vivo* PAM screening, guanine in the 2nd and the 3rd positions was indispensable for cleavage. However, in contrast to the tolerance for any nucleotide in the 4th position as disclosed *in vivo*, discrimination against cytosine was evidenced. Moreover, even though no specific nucleotides in the remaining positions were required for EHCas9 activity, a preference for thymine in the 5th position was revealed, suggesting the absence of this nucleotide in the *in vivo* screening might compromise target recognition when cytosine is present in the 4th position. In summary, while the PAMs compatible with EHCas9 target cleavage under the *in vitro* conditions used conformed to the consensus sequence 5’-NGGNNNN-3’, the optimal PAM seems to be 5’-NGGDTNN-3’ (D = A or T or G).

Next, we sought to further evaluate cytosine tolerance in the 4th position of the PAM along with the requirement of thymine in the 5th position. To this end, we carried out plasmid transformation assays equivalent to those used for *in vivo* PAM screening, but with individual plasmids instead of a PAM library containing in this case the target sequence adjacent to either 5’-TGGCG-3’, 5’-TGGTG-3’, 5’-TGGAG-3’, or 5’-TGGGG-3’ (Table S5). The motif 5’-TACCG-3’ was also surveyed as a PAM-negative control. As expected, when the target plasmid with the flanking 5’-TACCG-3’ sequence was transformed to cells expressing the three EH CRISPR-Cas components, the efficiency of transformation did not differ significantly from the efficiency observed in the absence of EHCas9. However, a marked decrease in the efficiency of transformation was found when 5’-TGGNG-3’ plasmids were transformed to EHCas9-expressing cells compared to hosts without the nuclease, exhibiting about four orders of magnitude difference in the case of the plasmid with cytosine in the 4th position of the PAM, and approximately five orders of magnitude for the rest (Fig. 3C). These results confirm that, even in the absence of thymine in the 5th position, EHCas9 efficiently elicits target cleavage in *E. coli* when any nucleotide is placed in the 4th position, being cytosine the one that sustains lower activity.

### Reaction conditions required for *in vitro* EHCas9-mediated target cleavage

To further characterize and refine the reaction parameters required for the double-strand break of DNA by EHCas9, *in vitro* cleavage assays were carried out under diverse conditions using an EH sgRNA and an 840-bp long linear DNA containing a target sequence. We opted for the 5’-TGGCG-3’ sequence at the place of PAM because it supported the weakest nuclease activity in the *in vivo* PAM validation experiments, serving us to maximize cleavage even for restrictive targets. Specific EH sgRNA-guided target cleavage will produce two dsDNA fragments (520-bp and 320-bp long, respectively). The 5’-TACCG-3’ sequence in the PAM region (PAM-negative target) was used as a negative control of cleavage (Table S5).

First, we assessed dsDNA target cleavage specificity at 37°C and the requirement for Mg^2+^ (Fig. 5A). To facilitate the formation of the ribonucleoprotein complex, we preincubated (15 min at 37°C) the nuclease with EH sgRNA (molar ratio 1:1) before mixing them with the target (the final Cas9:sgRNA:target molar ratio in the reaction solution was 20:20:1) in the presence of MgCl_2_. As expected, preincubation increased the rate of target cleavage when compared to reactions where all components were mixed simultaneously (*i.e*., 30 minutes after we added preincubated or non-preincubated protein and guide solutions to the target, 21.6% and 15.6% of the substrate was cleaved, respectively). Therefore, the subsequent *in vitro* experiments with EHCas9 and EH sgRNA were carried out after preincubation under the tested conditions. No cleavage products were noticed on the PAM-negative target, or when EH sgRNA or Mg^2+^ were not added to the reaction. In the presence of all the reactants, the substrate with the compatible PAM was cut once, generating only two DNA fragments whose sizes matched what was expected after cleavage within the target sequence. These results support that EHCas9 is an RNA-guided, sequence-specific, metal-dependent dsDNase.

**Fig. 5.**
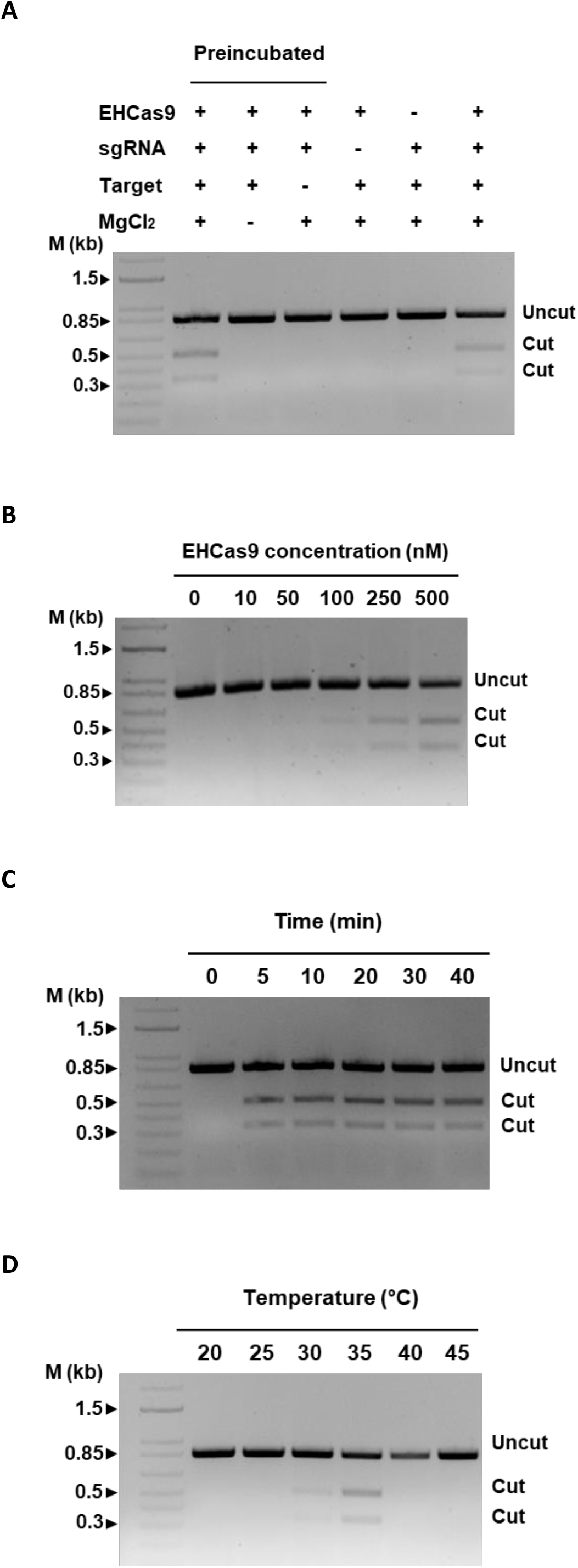
Representative agarose gel electrophoreses of *in vitro* EHCas9-mediated DNA cleavage reactions. Except indicated otherwise, experiments were performed under the following standard conditions: 30 min reaction time at 37°C, 20 mM MgCl_2_, 25 nM of an 840 bp linear DNA containing a target sequence and a compatible PAM, preincubated (15 min at 37°C) EHCas9:EH sgRNA mixtures (0.5 mM EHCas9 and 0.5 mM EH sgRNA concentration in the digestion reaction). The length of relevant bands (in kb) of a linear dsDNA molecular weight marker (M) and the position of bands corresponding to expected cut and uncut DNA substrates are indicated. (A) Samples from standard digestion reactions, mixed either after protein:guide preincubation (lane 2) or without preincubation (lane 7), and from reactions with a missing component (MgCl_2_, lane 3; a compatible PAM next to the target, lane 4; a guide, lane 5; the protein, lane 6) are included. (B) Samples from digestion reactions under standard conditions but for the protein concentration (up to 500 nM). (C) Samples from standard reactions incubated for up to 40 min. (D) Samples of digestions carried out under standard conditions except by the incubation temperature (from 20°C to 45 °C).

Next, we further characterized RNA-guided dsDNA cleavage activity in the presence of MgCl_2_, in terms of digestion time and temperature. To decide the amount of EHCas9 to be used in these experiments, constant concentrations of the EH sgRNA were preincubated for 15 min at 37°C with 10 nM to 0.5 μM of EHCas9 and subsequently mixed with a fixed concentration of substrate, so that the protein:sgRNA:substrate molar ratio in the digestion reaction varied from 1:50:2.5 to 20:20:1. Protein concentrations above 0.1 μM produced perceptible digestion products after 30 min, 0.5 μM EHCas9 being chosen for temperature and incubation time tests (Fig. 5B). When different reaction times (up to 40 min) were assessed at 37°C, even though a substantial proportion (21.6%) of the substrate was cut within the first 5 minutes underscoring the robustness of the nuclease, the maximum yield of cleavage observed (about 27% cleaved substrate) was reached after 30 min (Fig. 5C). Interestingly, incubation for 10 minutes longer did not increase the amount of substrate cleaved, suggesting that EHCas9 remains bound to the DNA after catalysing its cleavage, thus preventing it from acting on other target molecules. Regarding incubation temperature, cleavage products in digestion assays performed at 5°C intervals within the range 20-45°C were detected only at 30°C and 35°C, establishing a working temperature range of over 25°C to below 40°C, with optimum temperature around 35°C (Fig. 5D).

### EHCas9 tool is suitable for the positive selection of genome-edited *E. coli* cells

Next, we sought to investigate the applicability of EHCas9:sgRNA as a tool for the positive selection of *E. coli* mutants after recombineering assays (Fig. 6A). Two pBAD33-derivative plasmids (carrying a chloramphenicol resistance gene) were constructed, one with the genes encoding the two components of the tool (*i.e*., EHCas9 under the control of an arabinose inducible promoter and EH sgRNA under a constitutive promoter) and another plasmid to be used as a negative control of nuclease activity with just the EH sgRNA coding sequence. The EH sgRNA spacer matched a sequence of the chromosomal *pyrF* gene, located next to 5’-TGGAT-3’ in the PAM region Table S5). We co-transformed electrocompetent BW 27783 cells carrying the pKD46 plasmid expressing Lambda Red recombination proteins (Exo, Beta, Gam) [60] with the pBAD33 derivate carrying either the guide and *ehcas9* or just the guide (negative control), along with a 308 bp linear recombination template consisting of *pyrF* flanking sequences in such a way that recombination would lead to a 607 bp deletion within the gene. Transformants from three independent experiments were grown on chloramphenicol and arabinose containing plates and the *pyrF* region was PCR amplified from 90 randomly selected colonies (20 from each experiment with the plasmid expressing EHCas9 and 10 from each of the negative control replicas). Agarose gel electrophoresis of the PCR products invariably revealed a single band in each sample, running as a linear DNA fragment of size corresponding to either the deleted sequence in the case of clones expressing EHCas9 or the wild-type sequence for the negative control (Fig. 6B). These results demonstrate the robustness of our EHCas9 tool as a cell-killing agent and its suitability as a complement for applications benefiting from the positive selection of *E. coli* mutants, including genome editing.

**Fig. 6.**
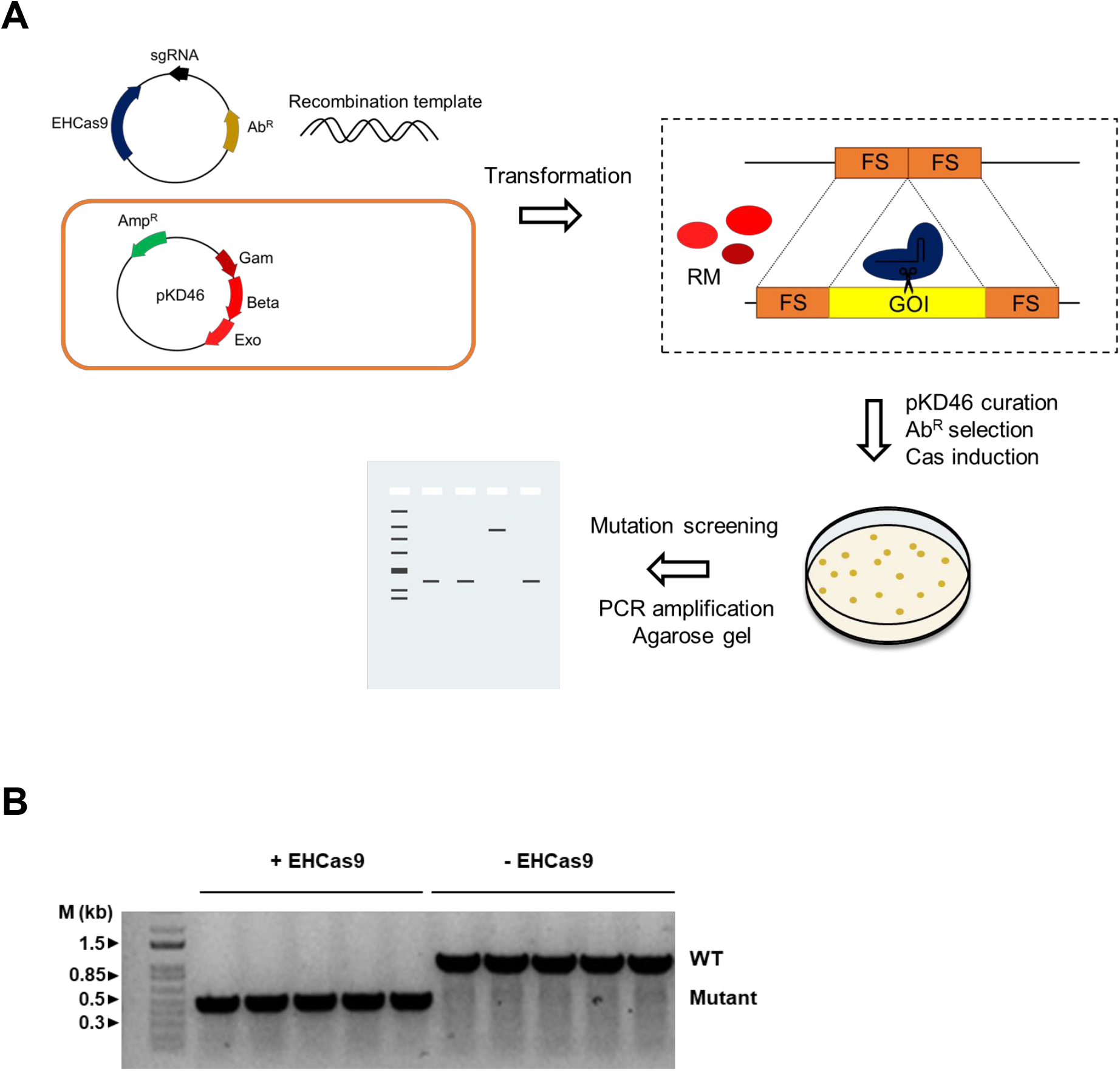
Prokaryotic genome editing with EHCas9. (A) Overview of the single-step general procedure for positive selection of genome-edited *E. coli. E. coli* cells harbouring a plasmid (*e.g*., pKD46; ampicillin resistance, Amp^R^) encoding Lambda Red recombination proteins (Gam, Beta, Exo) are co-transformed with an antibiotic resistance (Ab^R^) selectable plasmid carrying inducible *ehcas9* gene and a sgRNA targeting a sequence within the gene of interest (GOI) and a linear dsDNA template matching both sides of the target (flanking sites, FS). Recombination between the template and the flanking sites in the gene mediated by the Lambda Red recombination machinery (RM) will result in the deletion of the target sequence. The Cas9:sgRNA ribonucleoprotein complex will produce double-strand breaks in the non-edited target, often leading to cell death. Colonies grown at 37°C expressing Cas9 are selected on plates (*i.e*., arabinose and antibiotic-containing medium) and screened through PCR amplification and agarose gel electrophoresis to confirm the deletion. (B) An agarose gel electrophoresis of PCR products from the region surrounding *pyrF* (the gene of interest) in *E. coli* cells expressing the Lambda Red system from pKD46. DNA used for amplification was purified from chloramphenicol-resistant colonies grown in the presence of arabinose after co-transformation with a recombination template matching the flanks of *pyrF* (recombination would lead to a ca. 0.6 kb deletion), and a pBAD33-derivative plasmid (providing chloramphenicol resistance) encoding either both EHCas9 and an EH sgRNA that targets the *pyrF* gene (+ EHCas9) or only the EH sgRNA (-EHCas9). Each lane corresponds to a transformant clone. Bands running as linear DNA fragments with the length of the original (ca. 1 kb; WT) and the recombinant (ca. 0.5 kb; Mutant) *pyrF* region are indicated. The length of relevant bands of a linear dsDNA molecular weight marker is indicated.

### EHCas9 enhances genome editing in mammalian cells

Finally, the nuclease activity of EHCas9 was assessed in mammalian, N2a mouse cells, to evaluate its application as a genome editing tool in eukaryotes. The human codon-optimized *ehcas9* gene was chemically synthesized and inserted into a plasmid in such a way that the ORF was fused to an NLS coding region, being transcribed from the constitutive promoter CMV. EH sgRNA encoding sequence was inserted into another plasmid under the constitutive promoter U6 (Fig. 7A). The components of the SpCas9 genome editing tool cloned as the EH components were used as a reference.

**Fig. 7.**
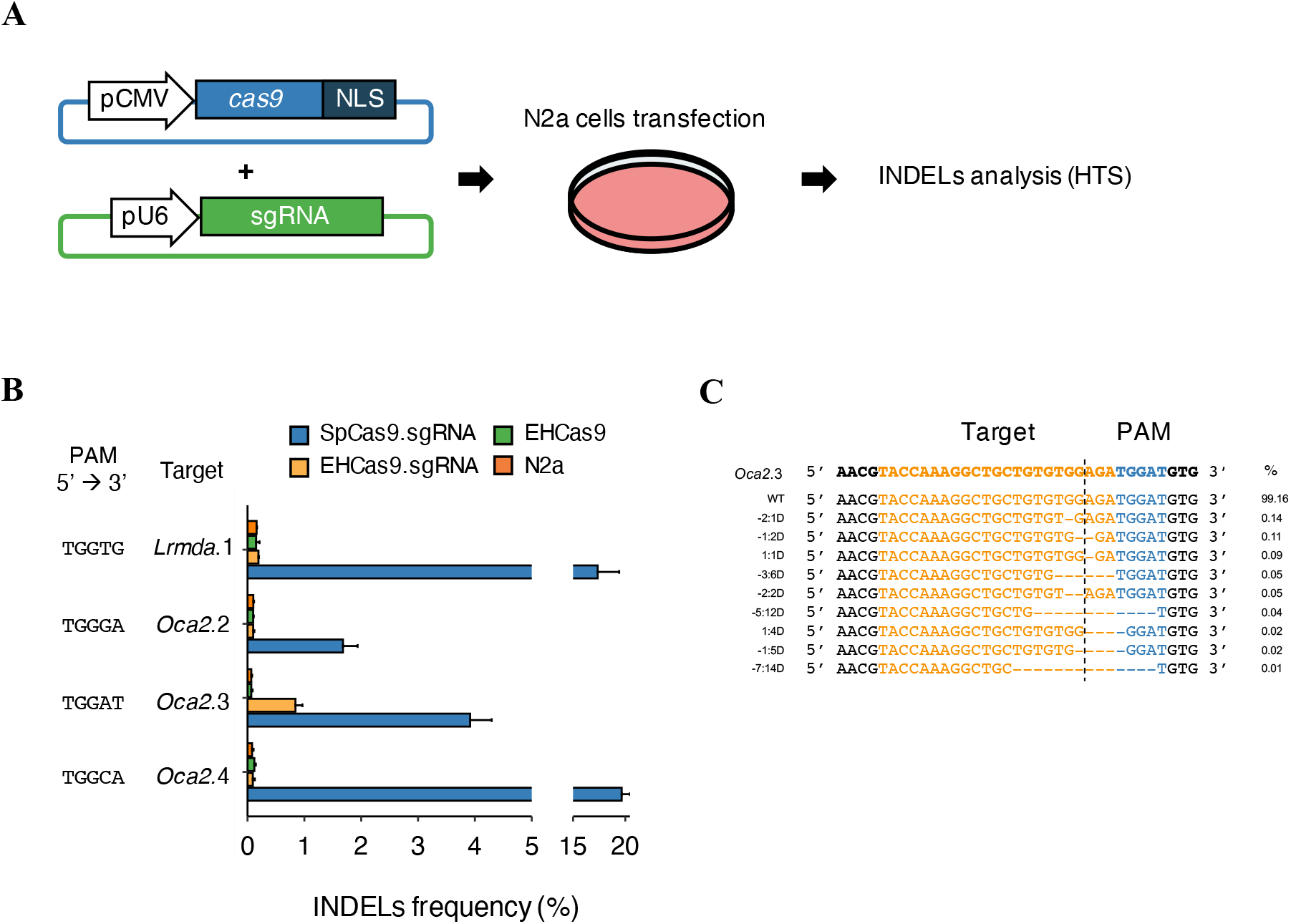
EHCas9-mediated genome editing in mouse cells. (A) Workflow of the Cas9 genome editing assay performed in mouse N2a cell cultures. A plasmid harbouring the *cas9* gene (*ehcas9* or *spcas9*) fused to a nuclear localization signal (NLS) was co-transfected into N2a cells with another plasmid carrying the corresponding sgRNA-encoding sequence under constitutive promoters (CMV and U6, respectively). Insertions and deletions (INDELs) in the guide complementary region were analysed by HTS of PCR amplicons from the region encompassing the target site. (B) Percentage of reads with INDELs (n=3 each, mean ± SD) obtained for 4 target regions (*Oca2.2, Oca2.3, Oca2.4, Lrmda.1*) after cell transfection with plasmids carrying SpCas9 and Sp sgRNA (SpCas9.sgRNA; blue), EHCas9 and EH sgRNA (EHCas9.sgRNA; yellow), or EHCas9 (EHCas9; green). Data from non-transfected cells (N2a; orange) are also included. The PAM sequence of each target site is indicated. (C) Alignment of the ten most abundant alleles revealed for *Oca2.3* in EHCas9:EH sgRNA experiments. Allele frequency (mean of three replicates) is displayed. Left column denotes deletion positions referred to the expected cleavage-site (dashed line) (*e.g*., −2:1D, one nucleotide deletion at position −2 from the cleavage site). Top line, original target region. The target sequence is coloured in orange and the PAM region in blue.

Four target regions of the mouse genome were tested, located in *Oca2* (*Oca2.2, Oca2.3, Oca2.4*) and *Lrmda* (*Lrmda*. 1) genes, adjacent to 5’-TGGGA-3’, 5’-TGGAT-3’, 5’-TGGCA-3’ and 5’-TGGTG-3’ in the PAM region, respectively (Fig. 7B and Table S5). The length of the sgRNA spacer region is a major determinant of the accuracy of target recognition [36,44,61–63]. We decided to use a 23-nt long spacer as this length is effective in most Cas9 proteins previously tested for mammalian genome editing, SpCas9 included. First, we evaluated the cell toxicity of the EHCas9 and SpCas9 tools by counting cell nuclei after transfecting N2a cells with plasmids encoding either the nuclease or the corresponding sgRNA. Even though a slight decrease in the number of nuclei relative to non-transfected cells was observed, no significant differences were found between the two Cas9 tools (Fig. S4). Therefore, this adverse effect on cellular growth was deemed acceptable for genome editing experiments. Then, we estimated EHCas9 performance as a genome editing tool by analysing insertion and deletion (INDEL) variations in the target site by HTS of the PCR-amplified target region after co-transfecting EHCas9 and EH sgRNA encoding plasmids in N2a cells (Fig. 7B). Negative controls lacking EH sgRNA were included and equivalent experiments were performed with components of the SpCas9 tool. Sequencing revealed INDELs for the four targets (conforming to the SpCas9 PAM 5’-NGG-3’) when challenged with the SpCas9 tool. However, in the case of EHCas9, INDELs were detected around the expected *Oca2.3* target site but not for the other targets. Remarkably, *Oca2*.3 was the only one where thymine was present in the 5th position of the PAM (5’-TGGAT-3’).

The efficiency of *Oca2*.3 editing was quantified as the proportion of reads with INDELs found in that sample, excluding other sequence variations that could be present in the population due to spontaneous mutations. The EHCas9 tool yielded 0.84 ± 0.12% reads with modified *Oca2*.3 sequence, whereas the editing efficiency found with SpCas9 was higher (3.92 ± 0.38% of the reads were edited). Still, the identity and relative frequency of the mutated alleles were similar for both proteins.

## Discussion

Our *in vitro* analyses of EHCas9 confirmed the typical biochemical features of Cas9 as a metal-dependent, RNA-guided dsDNase, that catalyses double-strand breaks at a fixed distance from compatible PAMs. Our results also suggested that EHCas9 has some properties, such as behaving as a single-turnover enzyme (like SpCas9 does) [64] and generating predominantly blunt-end cuts [6,13,22], that have been reported for most biochemically characterised orthologues, although multiple-turnover Cas9 (*i.e*., SauCas9) [22] and the generation of staggered termini [46] have also been observed. Still, staggered ends formation due to post-cleavage trimming or when targeting alternative sequences [22,46], as well as other here unexplored aspects exhibited by some orthologues such as RNA-independent DNA cleavage activity in the presence of Mn^2+^ ions [65], or nuclease activity on additional nucleic acids substrates (i.e., ssDNA, RNA, DNA:RNA hybrids) [66], cannot be dismissed. Additional experiments should be conducted to further characterize the enzyme in these aspects.

The selection of prokaryotic cells that have undergone mutations in genome editing experiments often requires laborious experimentation and Cas9 nucleases have been harnessed to simplify this task [26,59,67,68]. Here we have applied EHCas9 to positively select *E. coli* cells that had experienced deletion of a chromosomal target after recombinase-mediated homologous recombination, without requiring the introduction of sequences such as selection markers in the host genome. Remarkably, the sixty EHCas9-selected colonies that were examined carried the expected mutation. Moreover, in preliminary experiments (our unpublished results), transformation to *E. coli* of a plasmid carrying a chromosomal targeting guide and *ehcas9* encoding sequences produced only a few transformants which had depleted the *ehcas9* gene from the plasmid. These findings underline the cell toxicity of EHCas9 when guided to the *E. coli* chromosome and, therefore, its broad applicability as prokaryotic selection systems (*i.e*., sequence-specific antimicrobials) [30,69].

EHCas9 also proved active in mouse cells, enhancing genome editing of a target sequence. Compared to other Cas9-based tools that have been harnessed to applications in mammalian cells, EHCas9:sgRNA possesses several appealing features. First, EHCas9 size is in the range of the smallest orthologues (Table S6), which would permit eukaryotic cell delivery of the EHCas9 tool encoding components inserted into a single size-restricted vector, such as the adeno-associated viruses commonly used in biomedicine. Its size can also facilitate the administration of Cas9 derivates fused to peptides with distinct DNA-related activities, as has been done with dead-Cas9 (dCas9) proteins [70–74]. Second, being applicable in the context of biomedicine given its optimum operating temperature, its narrow working temperature range would allow for easy control of its activity: manipulation at room temperature would preclude digestion and a small temperature increase from the optimum temperature will terminate the reaction. Third, both in *E. coli* and *in vitro* experiments, EHCas9 cleaved dsDNA targets flanked by sequences in the PAM region matching the consensus 5’-NGGNNNN-3’, being guanine in the 2nd and the 3rd positions indispensable for cleavage as illustrated by their invariable occurrence in the top 10% most frequent PAMs identified in the *in vitro* PAM screenings. The requirement of such a short, frequently occurring motif, facilitates the selection of targets and its indispensability reduces the possibility of off-target effects on sequences that differ in any of the two positions. Still, a preferred consensus PAM 5’-NGGDT-3’ was inferred from the *in vivo* PAM validation assays and the *in vitro* PAM screening. Consistent with this prime PAM, in the mouse genome editing assays with the EHCas9 tool only the target flanked by 5’-TGGAT-3’ at the PAM region underwent INDELs. The absence of thymine in the 5th position (5’-TGGGA-3’, 5’-TGGCA-3’ and 5’-TGGTG-3’), or the presence of cytosine in the 4th position (5’-TGGCA-3’) of the PAM flanking the remaining targets probed, could account for the lack of INDELs detection in these regions when challenged with EHCas9. However, both cytosine in the 4th position and a nucleotide (*i.e*., guanine) other than thymine in the 5th position was tolerated for interference against target plasmids in *E. coli*, hinting at a lower performance of EHCas9 in mammalian cells. Previous studies with other Cas9 orthologs also reported similar discrepancies between hosts and targets as well as between *in vitro* and *in vivo* assessments [44,61,75]. Otherwise, in agreement with the more relaxed SpCas9 PAM (5’-NGG-3’), this tool produced mutations in the four targets tested in mouse cells, exhibiting a greater activity on the 5’-TGGAT-3’ associated target compared to EHCas9. The lower PAM tolerance and cleavage activity exhibited by EHCas9 results in a higher specificity relative to SpCas9. To confirm its specificity, it would be of great interest to assess EHCas9-mediated editing fidelity through genome-wide mutation analyses [76–78]. For applications in eukaryotic cells, EHCas9 will be particularly useful when a moderate activity on 5’-NGG-3’ linked targets is advisable, the tool further enabling the regulation of such an activity depending on the downstream sequence.

Metagenomic data mining has proved useful for the discovery of new CRISPR-Cas systems [47,48,79]. We detected the EH CRISPR-Cas in a metagenome obtained from a brackish-water (0.8% salinity) sample of a previously unexplored natural environment. To our knowledge, EHCas9 is the first type II CRISPR-Cas tool deployed from uncultivated microorganisms. Even though the DNA was purified from a subcellular fraction enriched in viruses, the limited genetic information available in the contig encoding the system prevented us from inferring the identity of the host genome. Regardless of its origin, the tool functions in heterologous hosts, both bacteria and mammalian cells, under physiological cell parameters that, allegedly, substantially differ from those faced by the natural carrier. The performance of this tool highlights the potential of metagenomic analysis of environments, including those distantly related to the intended recipient, to expand the CRISPR toolbox.

## Materials and methods

### Generation of metagenomic data

Water samples collected from a lagoon in El Hondo Natural Park (Spain) were first pre-filtered through filter paper and a 5 μm pore size Durapore® membrane filter (Merk). Subsequently, sequential filtration through a 0.22 μm pore size Durapore® membrane filter (Merk), and a VIVAFLOW 200 30,000 MWCO (Sartorius) crossflow ultrafiltration device was performed. The filtered sample was further concentrated using a 3K Ultra Amicon® Centrifugal filter (Millipore). DNA was purified from the concentrate with PureLink® Viral RNA/DNA Mini kit (Invitrogen).

DNA sequencing was performed by FISABIO-Public Health using Illumina HiSeq with paired-end 150 cycles. Low-quality reads were trimmed with the PRINSEQ-lite program [80] using the settings: min_length: 50, trim_qual_right: 30, trim_qual_type: mean, and trim_qual_window: 20. Then, eukaryotic sequences were identified with BLASTn searches (options: -taxidlist: taxid:2759, -evalue: 0.005), against the NCBI non-redundant nt database (https://blast.ncbi.nlm.nih.gov/Blast.cgi). Read sequences with an identity (number of identical positions/query length) > 0.9 were filtered out from the total reads using the FastQ.filter.pl script of the Enveomics Collection [81]. De novo assembly of the remaining reads was conducted with SPAdes v3.13.0 program [82] using metaspades option with parameters: -k 21, 33, 55, 77, 99, 127.

### Computational analysis to detect type II CRISPR-*cas loci* in the metagenomic dataset

For the identification of CRISPR-Cas systems in our metagenome, ≥ 2 kb-long scaffolds were first enquired with the CRISPRCasFinder (CCFinder) standalone program to find *cas* genes and CRISPR arrays [83]. Next, open reading frames (ORFs) from the 745 contigs containing CRISPR-Cas components thus identified were predicted using Prodigal v2.6.3 [84] and the derived catalogue of protein sequences was queried with the Cas9 Hidden Markov Model (HMM) protein domain profiles (Table S1) using the hmmersearch program of the HMMER package v3.2 [85,86], followed by manual data curation.

As a first step for the identification of putative tracrRNA-encoding regions, degenerated repeat sequences were searched in the vicinity of the CRISPR-*cas* loci with the Benchling online platform (https://benchling.com/editor). Then, promoter and terminator sequences were predicted on either side of the repeat-matching regions with BPROM and FindTerm [86], respectively. A system (EH CRISPR-Cas system) associated with a *cas9* gene (*ehcas9*) and a putative tracrRNA was selected for further analysis.

### Phylogenetic analysis and EH sgRNA design

A phylogenetic tree of EHCas9 protein was constructed using bioinformatic tools available at the NGPhylogeny.fr website (https://ngphylogeny.fr) [87]. Specifically, a multiple protein sequence alignment of EHCas9 with a Cas9 orthologues dataset compiled by Gasiunas and collaborators [46], was carried out with the MUSCLE software [88]. The tree was generated from the alignment with the Fast Tree program using a JTT evolutionary model and discrete gamma model [89,90].

An EH sgRNA was inferred after the alignment, using CLUSTAL Omega [91], of EH CRISPR repeat and the predicted EH tracrRNA sequences with the corresponding elements from *Sulfitobacter donghicola*. The secondary structure of the EH sgRNA was predicted with the Mfold program (http://www.unafold.org) [92].

### Protein structure prediction and analysis

EHCas9 3D structure was modelled using ColabFold: AlphaFold2 using MMseqs2 [93,94]. The following configuration was set up: pdb70 template_mode; MMseqs2 (UniRef+Environmental) msa_mode; unpaired+paired pair_mode; other options by default. After this initial analysis, the list of templates used was checked and structures corresponding only to resolved Cas9-sgRNA-DNA ternary complexes (5axw, 5czz, 5×2g, 5×2h, 5y36, 6jdv, 6joo, 6kc8, 6m0w, 6m0x, 6wbr and 7el1) were subjected to a second prediction, setting up template_mode as custom. PyMOL program (https://github.com/schrodinger/pymol-open-source) was used to visualize, represent and analyse de resulting models.

### Bacterial strains, growth conditions, oligonucleotides and plasmids used

Bacterial strains and plasmids used in this work are listed in Table S2 and Table S3, respectively. Oligonucleotides were purchased from IDT and are listed in Table S4.

Unless otherwise specified, *Escherichia coli* cultures were grown at 37°C in Luria-Bertani (LB) broth under rotational shaking at 180 rpm or on LB agar. For the selection of plasmid carriers, media were supplemented with chloramphenicol (25 μg/ml), ampicillin (100 μg/ml), spectinomycin (50 μg/ml) or kanamycin (50 μg/ml), as appropriate.

### Plasmid cloning methods and transformation procedures

Guide spacer sequences were inserted into pMML13 plasmid using the Golden Gate method [95]. The other molecular cloning trials and plasmid gene replacements were performed by Gibson’s assembly with Gibson Assembly® Cloning Kit (NEB). The procedure used to generate each plasmid construct is described in the corresponding subsection of Material and Methods.

For the preparation of electrocompetent *E. coli* BL21(DE3) and *E. coli* BW27783 cells, stationary-phase liquid cultures were diluted 1/100 in sterile LB broth and grown to OD_600_ = 0.5. Cells collected by centrifugation were washed three times in deionized water and once in 10% glycerol. Transformations were performed with 50 μl of freshly prepared electrocompetent cell suspensions incubated on ice for 25 min after adding the DNA. The cells:DNA mixture was transferred to an ice-cold 2 mm gap electroporation cuvette (Molecular Bioproducts) and electroporated at 2.5 kV with a MicroPulser (BIORAD). One ml of Super Optimal broth with Catabolite repression (SOC) was immediately added to the cell suspension and the entire volume was incubated for 1 hour under standard growth conditions in a 12 ml tube. Finally, cells were plated to media supplemented with the corresponding antibiotic for plasmid selection and incubated overnight at either 30°C in the case of the thermosensitive plasmid pKD46 or at 37°C in all other cases.

Chemically competent *E. coli* NZYStar (NZYTech) and *E. coli* TOP10 (Invitrogen) cells were transformed as indicated by the manufacturer.

### DNA purification and analysis

Plasmids were isolated from *E. coli* with either PureLink™ HiPure Plasmid Midiprep Kit (Invitrogen) or PureLink™ HiPure Plasmid Miniprep Kit (Invitrogen). PCR products and DNA fragments were purified with GFX™ PCR DNA and Gel Band Purification Kit (Cytiva).

The concentration and purity of nucleic acid solutions were estimated with NanoDrop ND-1000 Spectrophotometer (Thermo Scientific), and their integrity was assessed by agarose gel electrophoresis.

To visualize agarose gel electrophoresed DNA molecules, gels containing GreenSafe premium (NZYTech) nucleic acid stain were imaged with ChemiDoc XRS+ Gel Imaging System (BIORAD). The 1 Kb Plus DNA Ladder (Invitrogen) was included in the agarose gels as a DNA weight marker.

### *In vivo* PAM screening and validation

For *in vivo* EHCas9 PAM screening, the pMML01 plasmid to be used as a negative control of EHCas9 activity, was generated by inserting an EH CRISPR array made of two 36-bp long repeats separated by a 29-bp long spacer in pBAD33. We engineered another pBAD33-derivative plasmid (pMML02) bearing the *ehcas9* gene as well, and a pUC57-based plasmid (pMML03) carrying a 300-bp long insert encompassing the predicted EH tracrRNA coding sequence. To construct pMML02, an *E. coli* codon-optimized *ehcas9* gene and the CRISPR array flanked by a constitutive promoter (Part:BBa_J23101, BioBricks collection) and the terminator sequence BBa_B1006, purchased as G-blocks from NZYTech, were inserted into pMML01 in such a way that *ehcas9* was under the control of the P_BAD_ arabinose promoter. For the construction of pMML03, the EH CRISPR-Cas intergenic regions synthesized as a G-block by NZYtech were cloned under the *lac* promoter of pUC57. A 3-nt PAM randomized, pSEVA431-derivative (spectinomycin resistance) plasmid library was generated by PCR mutagenesis with primers containing random nucleotides (Table S4) at the 2nd, the 3rd, and the 4th positions 3’ from the 5’-CCTGTATATCGTGCGAAAAAGGATGGATA-3’ target sequence in the spacer-matching strand (Table S5).

Electrocompetent *E. coli* BW 27783 cells were co-transformed with pMML03 and either pMML01 or pMML02 and selected on ampicillin and chloramphenicol containing LB agar plates. Transformant colonies were grown on LB broth supplemented with ampicillin, chloramphenicol, 0.2% L-arabinose and 1mM IPTG. Then, electrocompetent cells were prepared from cultures at OD_600_ = 0.5 and three independent transformation experiments with 300 ng of the PAM library were performed for both pMML01 and pMML02 carriers. Transformants carrying pSEVA431-derivative plasmids were selected on LB agar supplemented with spectinomycin and plasmids were isolated from ca. 10^5^ pooled colonies. The plasmid region surrounding the PAM was PCR amplified with barcode-containing oligonucleotides (see Table S4 for further details) and sequenced by High Throughput Sequencing (HTS) using an Illumina NovaSeq PE250 sequencing system (Novagene). The proportion of reads with each specific PAM sequence obtained from pMML02 carriers was compared with the corresponding values from cells carrying the negative control pMML01 to estimate their log2 fold-change. PAM sequences with a log2 value higher than 7 were used to generate sequence logos with the WebLogo application (https://weblogo.berkeley.edu/logo.cgi).

A subset of PAM sequences was validated for EHCas9 activity by similar interference assays except for the transformation with 200 ng of individual pSEVA431-derivative plasmids pMML04 - pMML07 carrying specific PAM sequences, instead of with a plasmid library. These plasmids were generated by PCR mutagenesis using primers with different nucleotides at the 2nd, the 3rd, and the 4th positions (Table S4). The efficiency of transformation was estimated as the number of transformant colony forming units (CFU) per μg of DNA.

### *In vitro* PAM screening and cleavage site identification

The sequences in the PAM region compatible with dsDNA target cleavage by EH sgRNA-guided EHCas9 were assessed *in vitro* following a previously described procedure [46]. First, *in vitro* translated EHCas9 at 0.5 μM final concentration was mixed with 2 μg of *in vitro* transcribed sgRNA to facilitate binary nucleoprotein complex formation. After 15 minutes at room temperature, 1 μg of a 7-bp PAM randomized target plasmid library was added to 10 μl of the complex solution and incubated at 37°C in 100 μl reaction buffer containing 10 mM Tris-HCl pH 7.5, 100 mM NaCl, 10 mM MgCl_2_, 1 mM 1,4-dithiothreitol (DTT). After one hour of incubation, adapters were ligated to the ends of the linear DNA fragments generated by plasmid cleavage. Finally, these fragments were PCR amplified and sequenced using MiSeq Personal Sequencer (Illumina). The occurrence of PAM sequences found in these reads (treatment frequency) was normalized to that found in equivalent control experiments conducted in the absence of EHCas9, according to the equation: (treatment frequency) x (average control frequency) / (control frequency).

PAM preferences were inferred from the normalized occurrence of each nucleotide at each position utilizing position frequency matrices. Only the top 10% of most frequent PAMs were considered for WebLogo generation. The cleavage site was inferred from the most frequent read termini at the target region.

### Recombinant protein purification

For heterologous expression of EHCas9 protein, *E. coli* codon-optimized *ehcas9* gene supplied by NZYtech was fused to an N-terminal hexahistidine tag under a *lac*/IPTG-inducible promoter in a pHTP1 vector, generating pMML22 plasmid. *E. coli* BL21(DE3) previously transformed with pMML22 was grown at 37°C in LB broth supplemented with kanamycin. When the culture reached OD_600_ = 0.5, protein expression was induced by adding 1mM IPTG and after 16 h of incubation at 16°C cells were harvested by centrifugation (5.000 x g for 15 min at 4°C) and resuspended in binding buffer composed of 50 mM pH 7.6 phosphate buffer, 500 mM NaCl, 10 mM imidazole, 5% glycerol, 10 mM ß-mercaptoethanol, 1mM phenylmethylsulfonyl fluoride (PMSF). Cells were disrupted by sonication with a Branson Digital Sonifier®. After centrifugation (23.700 x g for 25 min at 4°C), the supernatant was loaded on a 1 ml HisTrap HP column (GE Healthcare), washed with 20 volumes of binding buffer, and eluted with elution buffer (50 mM pH 7.6 phosphate buffer, 500 mM NaCl, 150 mM imidazole, 5% glycerol, 10 mM ß-mercaptoethanol, 1 mM PMFS). The eluted fraction was concentrated to 1 ml in cleavage buffer (50 mM pH 7.6 phosphate buffer, 150 mM NaCl, 5% glycerol, 10 mM ß-mercaptoethanol) using Amicon Ultra centrifugal filters (Millipore) and loaded on a gel filtration HiLoad™ 16/600 Superdex™ 200 pg (Cytiva). Elution fractions were analysed through SDS-PAGE and the one containing a protein of the size expected for EHCas9 was concentrated as above (Fig. S1).

NZYBlue Protein Marker (NZYtech) was used for protein size estimation and protein concentrations were measured with QUBIT® 2.0 (Invitrogen).

### Optimization of EHCas9-mediated DNA cleavage assays

For the optimization of the reaction conditions required by EHCas9 for dsDNA cleavage, an *in vitro* transcribed EH sgRNA was generated. To that extent, oligonucleotides carrying a T7 promoter and a 23-nt long spacer-matching sequence in pSEVA431 (see Table S4 for further details) were used to obtain a dsDNA template by PCR amplification of the constant sgRNA-encoding region from the pMML08 plasmid. The amplicon was transcribed with HiScribe T7 Quick (NEB) following the manufacturer’s instructions, including the optional DNase treatment, and the RNA was purified with Monarch® RNA clean-up kit (NEB). sgRNA aliquots were stored at −80°C.

An 840-bp PCR fragment containing a target with 5’-TGGCG-3’ PAM was amplified from pMML05 (Table S3) and used as a cleavage substrate. As a control, a similar fragment was amplified from pMML05 containing a target with 5’-TACCG-3’.

Unless otherwise stated, cleavage assays were performed at 37°C for 30 min in a 50 μl reaction buffer containing 20 mM MgCl_2_, 25 μM DNA substrate and 0.5 μM EHCas9 premixed with EH sgRNA (50 μM final concentration) for 15 min at 37°C. Digestion was quenched on ice and DNA purified as above indicated. Reaction products were analysed by electrophoresis in 1% agarose gels. The intensity of the bands was estimated with Image Lab software (BIO-RAD) and cleavage efficiency was calculated for each sample as a ratio of the intensity of the bands corresponding to cleaved products, to the sum of the intensities of all the bands in the lane.

### Positive selection of *E. coli* mutants

For the selection of genome-edited *E. coli* cells, the pMML09 plasmid encoding EHCas9 and a sgRNA targeting the chromosomal *pyrF* gene was constructed from pMML02 by replacing the region between the promoter and terminator of the CRISPR array with a sgRNA-coding sequence containing a *pyrF*-matching spacer.

As a negative control of EHCas9 activity, an *ehcas9*-depleted plasmid (pMML10) was generated through PCR amplification of pMML09.

A linear DNA recombination template comprising a 145-bp sequence matching the intergenic upstream region of *pyrF* and a 163-bp sequence matching the downstream region of the gene was generated by Gibson assembly.

Electrocompetent *E. coli* BW 27783 cells were transformed with pKD46 (ampicillin resistance) encoding the Lambda Red recombination system [60]. As replication of this plasmid is temperature-sensitive, being cured at 37°C, transformants were grown on ampicillin containing LB agar plates incubated at 30°C. pKD46 carrier colonies were transferred to LB broth supplemented with ampicillin and grown at 30°C to OD_600_ = 0.2. Then, 0.2% L-arabinose was added to induce expression of the Lamba Red proteins and electrocompetent cells were prepared from the culture when OD_600_ = 0.5 was reached. Next, 3 aliquots were co-transformed with 150 ng of DNA template and 50 ng of either pMML09 or pMML10. Transformant colonies were grown at 37°C, preventing pKD46 replication, on LB agar supplemented with chloramphenicol (pMML09 and pMML10 plasmid selection) and 0.2% L-arabinose (induction of *ehcas9* transcription). *pyrF* integrity was assessed by 1% agarose gel electrophoresis of the PCR amplified genomic region.

### Genome editing of mammalian cells

For genome editing assays, hCas9 plasmid (Addgene #41815; Mali *et al*. 2013) carrying *spcas9* gene fused to a nuclear location sequence (NLS) tag controlled by a constitutive cytomegalovirus (CMV) promoter, and MLM3636 plasmid (Addgene #43860) encoding a compatible sgRNA (Sp sgRNA) backbone under the constitutive U6 promoter, were used as scaffolds to construct equivalent plasmids where the SpCas9 and Sp sgRNA encoding sequences were replaced with the human codon-optimized *ehcas9* gene (pMML12) and an EH sgRNA constant region (pMML13), respectively. The two inserts were purchased from NZYTech as G-blocks.

sgRNA targets within *Oca2* and *Lrmda* genes of the mouse genome (Table S5) were selected with Breaking–Cas software (http://bioinfogp.cnb.csic.es/tools/breakingcas/) [96] using the search parameters: *PAM sequence:* TGGN, *PAM position:* 3’, *guide length:* 23 nts, *mismatches:* Up to 4. sgRNA spacer regions matching the selected targets were inserted into pMML13 and MLM3636 to generate plasmids pMML14-pMML17 and pMML18-pMML21, respectively.

Neuro-2a (N2a) cells of *Mus musculus* (mouse neuroblasts; ATCC, CLC-131 ™) were maintained in Dulbecco’s Modified Eagle’s Medium (DMEM) – high glucose (Sigma) supplemented with sterile-filtered 10% fetal bovine serum, 10 mM HEPES pH 7.4, 2 mM L-glutamine, 100 IU/ml penicillin and 100 μg /ml streptomycin, at 37°C with 5% CO_2_ and 95% humidity.

For toxicity assessment of CRISPR-Cas components, cells were plated in 96-well plates at a density of 1.5·10^4^ cells/mL per well in a total volume of 100 μl of DMEM without antibiotics and co-transfected with 200, 150 and 100 ng of pMML12 or hCas9 and 100 ng of pMML13 or MLM3636, respectively. Three days after transfection, cells were fixed with 4% paraformaldehyde for 30 min at room temperature and cell nuclei were DAPI stained and counted with a fluorescent microplate multimode reader Spark® (TECAN).

For *in vivo* genome editing experiments, N2a cells were plated in 24-well plates at a density of 4·10^5^ cells/mL per well in a total volume of 500 μl of DMEM without antibiotics and co-transfected with 1 μg of pMML12 or hCas9 and 500 ng of the corresponding sgRNA-encoding plasmid (pMML18 – pMML21 or, pMML14 – pMML17 correspondingly). Genomic DNA was extracted with the High Pure PCR Template Preparation kit (Roche) from cells collected 72 hours after transfection.

Transfections were carried out with Lipofectamine 2000 (Invitrogen), following the manufacturer’s instructions.

### INDELs frequency analysis

The results of mouse genome editing were analysed by insertions and deletions (INDELs) formation [97]in the target sites. To this end, 300-400 bp amplicons were generated by PCR amplification of the regions surrounding the target, using 100 ng of N2a genomic DNA as a template. PCR products were sequenced at Novogene using Illumina NovaSeq 6000 with paired-end 250 cycles. Low-quality reads and adaptors were trimmed with Trimmomatic v0.39 (parameters: java –jar trimmomatic-0.39.jar PE ILLUMINACLIP:2:30:10 SLIDINGWINDOW:4:15 MINLEN:50) [97]. Sequencing reads were mapped to the reference target sequence with the Bowtie2 v2.4.2 program [98] and converted to BAM files format with the Samtools package [99]. The analysis of INDELs was carried out with R Core Team (2021) using CrispRVariants 1.20.0 package [100].

## Accession number

The sequence of the metagenomic contig encoding the EH CRISPR-Cas system is deposited in GenBank with accession number OP485146.

## Acknowledgements

We are thankful to Riccardo Rosselli, Manuel Martínez.García and Noemi M. Guzmán (Universidad de Alicante) for valuable input.

## Supplemental material

Fig. S1. SDS polyacrylamide gel electrophoresis showing the steps of His-tagged EHCas9 purification. A whole lysate of bacteria expressing EHCas9 (Lysate) and samples of protein extracts purified through His-binding column (His-column) as well as after subsequent gel filtration (Gel filtration) are included. The size of bands corresponding to the protein molecular weight marker (M) is indicated. The major band of the protein extracts corresponds to a protein around 120 kDa as expected for the His-tagged EHCas9.

Fig. S2. Amino acid sequence alignment of Cas9 proteins. (A) Pairwise sequence alignment of EHCas9 with the closest structurally characterized orthologue from Corynebacterium diphtheriae (CdCas9; protein data bank ID: 6JOO). The boundaries of RuvC (RuvCI-III motifs), Bridge Helix (BH), recognition (REC), HNH, Phosphate Lock Loop (PLL), WED, and PAM-Interacting (PI) domains of CdCas9 are denoted by colored bars bellow the sequence. (B) Multiple sequence alignment of EHCas9 with structurally characterized orthologues: CjCas9, Campylobacter jejuni; NmCas9, Neisseria meningitidis 8013; StCas9, Streptococcus thermophilus LMD9; SaCas9, Staphylococcus aureus; SpCas9, Streptococcus pyogenes. Some of the EHCas9 amino acid positions are listed. The RuvC and HNH catalytic sites are red and blue shaded, respectively. In both panels, conserved positions are marked with an asterisk.

Fig. S3. EHCas9 predicted structures. (A) Cartoon representation of the EHCas9 3D structure predicted by AlphaFold2. α-helixes, β-sheets and loops are depicted. (B) Surface representation of the EHCas9 3D structure predicted by AlphaFold2. Protein domains are colored according to the legend. The typical bi-lobed structure of Cas9 can be observed. (C) Cristal structure of CdCas9 PI domain. (D) PI domain structure of EHCas9 predicted by AlphaFold2. The conserved core fold formed by antiparallel β-sheets that participate in PAM recognition can be observed.

Fig. S4. Growth of N2a mouse cells expressing components of the EH and SpCas9 tools. (A) Total counts of nucleated cells in non-transfected cells (No plasmid) and after transfection with 200 ng, 150 ng and 100 ng of plasmids encoding SpCas9 or EHCas9. (B) Total counts of nuclei in non-transfected cells (No plasmid) and after transfection with 100 ng of plasmids encoding sgRNA of either SpCas9 (Sp sgRNA) or EHCas9 (EH sgRNA) (n=3, mean ± SD). (C) DAPI staining of non-transfected N2a cells (N2a) and cells after transfection with plasmids encoding SpCas9 or EHCas9.

Table S1. Hidden Markov Models used in this work for the identification of Cas9 proteins in the metagenome dataset.

Table S2. *E. coli* strains used in this work.

Table S3. Plasmids used in this work.

Table S4. Oligonucleotides used in this work.

Table S5. Cas9 target sequences used in this work.

Table S6. Relevant features of native Cas9 orthologues applied to genome editing in mammalian cells.

